# Biomarkers of Memory Variability in Traumatic Brain Injury

**DOI:** 10.1101/2020.07.27.223073

**Authors:** Richard Adamovich-Zeitlin, Paul A. Wanda, Ethan Solomon, Tung Phan, Bradley Lega, Barbara C. Jobst, Robert E. Gross, Kan Ding, Ramon Diaz-Arrastia, Michael J. Kahana

## Abstract

Traumatic brain injury (TBI) is a leading cause of cognitive disability and is often associated with significant impairment in episodic memory. In TBI survivors, as in healthy controls, there is marked variability between individuals in memory ability. Using recordings from indwelling electrodes, we characterized and compared the oscillatory biomarkers of mnemonic variability in two cohorts of epilepsy patients: a group with a history of moderate-to-severe TBI (*n* = 37) and a group of non-TBI controls (*n* = 111) closely matched for demographics and electrode coverage. Analysis of these recordings demonstrated that increased high frequency power and decreased theta power across a broad set of brain regions mark periods of successful memory formation in both groups. As features in a logistic-regression classifier, spectral power biomarkers effectively predicted recall probability, with little difference between TBI and non-TBI controls. The two groups also displayed similar patterns of theta-frequency connectivity during successful encoding periods. These biomarkers of successful memory, highly conserved between TBI patients and controls, could serve as the basis for novel therapies that target disordered memory across diverse forms of neurological disease.

## Introduction

Traumatic brain injury (TBI) produces lasting impairments in episodic memory and executive function. Neuropsychological tests reveal some of the most profound functional impairments in measures of delayed recall, a particularly sensitive index of episodic memory function (Dikmen *et al*., 2009; Vakil, in press). Cognitive rehabilitation therapy has been the primary treatment strategy for patients suffering from memory and executive dysfunction, but therapeutic evidence has been mixed, leading to continued uncertainty over clinical best practices (Shoulson *et al*., 2012), and no medication constitutes a clinical standard to address cognitive impairment in the TBI population (Cicerone *et al*., 2006). To develop effective therapies for treating such memory loss, we must first understand whether and how the physiology of memory function differs between TBI-affected individuals and their non-TBI counterparts.

TBI is a heterogeneous injury, resulting in both focal and diffuse pathologies, and the mechanisms of memory dysfunction may be distinct from those resulting from developmental or neurodegenerative conditions. Post-traumatic axonal degeneration may contribute to hippocam- pal atrophy, reduced volume of white matter, and passive ventricular expansion (Bigler and Maxwell, 2011). Structural and functional neuroimaging studies in patients with moderate-to-severe TBI have highlighted white matter lesions (Smith *et al*., 2003) and reduced connectivity among the frontal, temporal, and parietal lobes (Wang *et al*., 2011; Pandit *et al*., 2013). This pattern of damage, known as diffuse axonal injury, may underlie slowed cognitive processing (Spitz *et al*., 2013).

Although neuroimaging studies provide an important first step towards understanding the changes in brain structure and networks in TBI, they do not provide critical information as to whether or how TBI alters the electrical activity underlying affected cognitive processes such as episodic memory. Indeed, research remains ongoing regarding the effectiveness of EEG as a diagnostic tool for brain injury (Nuwer *et al*., 2005; Slobounov *et al*., 2012; Edlow *et al*., 2017). Given the critical importance of electrophysiology for tracking the temporal dynamics of memory function, and the potential of using such biomarkers to control neuromodulatory therapy (Ezzyat *et al*., 2018; Hanslmayr *et al*., 2019), we sought to address this gap in our knowledge by investigating the neural correlates of episodic memory using direct brain recordings in epileptic patients with and without a history of moderate-to-severe TBI.

Previous work by multiple research groups has revealed a clear set of physiological biomarkers of intraindividual variability in human memory function. Specifically, it is now well-established that high-frequency activity (generally *>* 30 Hz) increases across a broad network of brain regions during periods of successful memory encoding and retrieval (see (Burke *et al*., 2015) for a review). Similarly, increases in theta band (3-8 Hz) functional connectivity, accompanied by decreases in theta band power, mark periods of successful encoding and retrieval in both the medial temporal lobe (Solomon *et al*., 2019; Hanslmayr *et al*., in press) and neocortex (Burke *et al*., 2013; Solomon *et al*., 2017).

The present study aims to determine whether these biomarkers of variability in memory encoding and retrieval are conserved across TBI and non-TBI cohorts. As direct brain recordings may only be ethically obtained in human volunteers undergoing intracranial EEG for indications such as intractable epilepsy, and because these patients occasionally have significant prior history of TBI (a known risk factor in developing epilepsy (Verellen and Cavazos, 2010; Ding *et al*., 2016)), we conducted a careful chart review of more than 300 neurosurgical patients who performed memory testing as part of a multi-center memory study. We identified 37 patients with a history of moderate-to-severe TBI and 111 matched controls. Although findings from this public data-set have been reported in prior publications, none have considered the potential relation between TBI history and biomarkers of successful memory.

## Materials and methods

We analyzed direct neural recordings from cortical and deep brain structures in patient-subjects while they performed verbal episodic memory tasks. These patients were surgically implanted with intra-parenchymal recording electrodes to directly measure intracranial electroencephalographic (EEG) activity. We quantified each subject’s neural biomarkers of successful memory encoding by analyzing two distinct features of the EEG time-series across all implanted electrodes during memory encoding: spectral power across a broad range of frequencies and phase-locking connectivity in the theta (3-8 Hz) band. We compared these features between the groups to discern if a history of moderate-to-severe traumatic brain injury was associated with a difference in the electrophysiology of successful memory encoding. We then asked if these features could be used to reliably predict successful memory encoding in these TBI patients.

### Human Subjects

We examined data from 148 neurosurgical patients (105 male, mean age = 41) with medication-resistant epilepsy undergoing invasive monitoring for seizure localization. Neurosurgeons implanted patients with intra-parenchymal depth electrodes and/or subdural grids and strips placed on the cortical surface. The location of these electrodes was determined based solely upon clinical considerations. As such, there are no two patients with the same electrode configuration, but when aggregated, the coverage is expansive across the entire brain **(Figure 1B**). The included data were collected as part of a larger project in collaboration with Columbia University Medical Center (New York, NY), Dartmouth-Hitchcock Medical Center (Hanover, NH), Emory University Hospital (Atlanta, Georgia), Hospital of the University of Pennsylvania (Philadelphia, PA), the Mayo Clinic (Rochester, MN), Thomas Jefferson University Hospital (Philadelphia, PA), the National Institute of Neurological Disorders and Stroke (Washington, D.C.), and University of Texas Southwestern Medical Center (Dallas, TX). Institutional review boards at the respective hospitals approved the research protocol, and we obtained informed consent from each patient.

**Figure 1.**
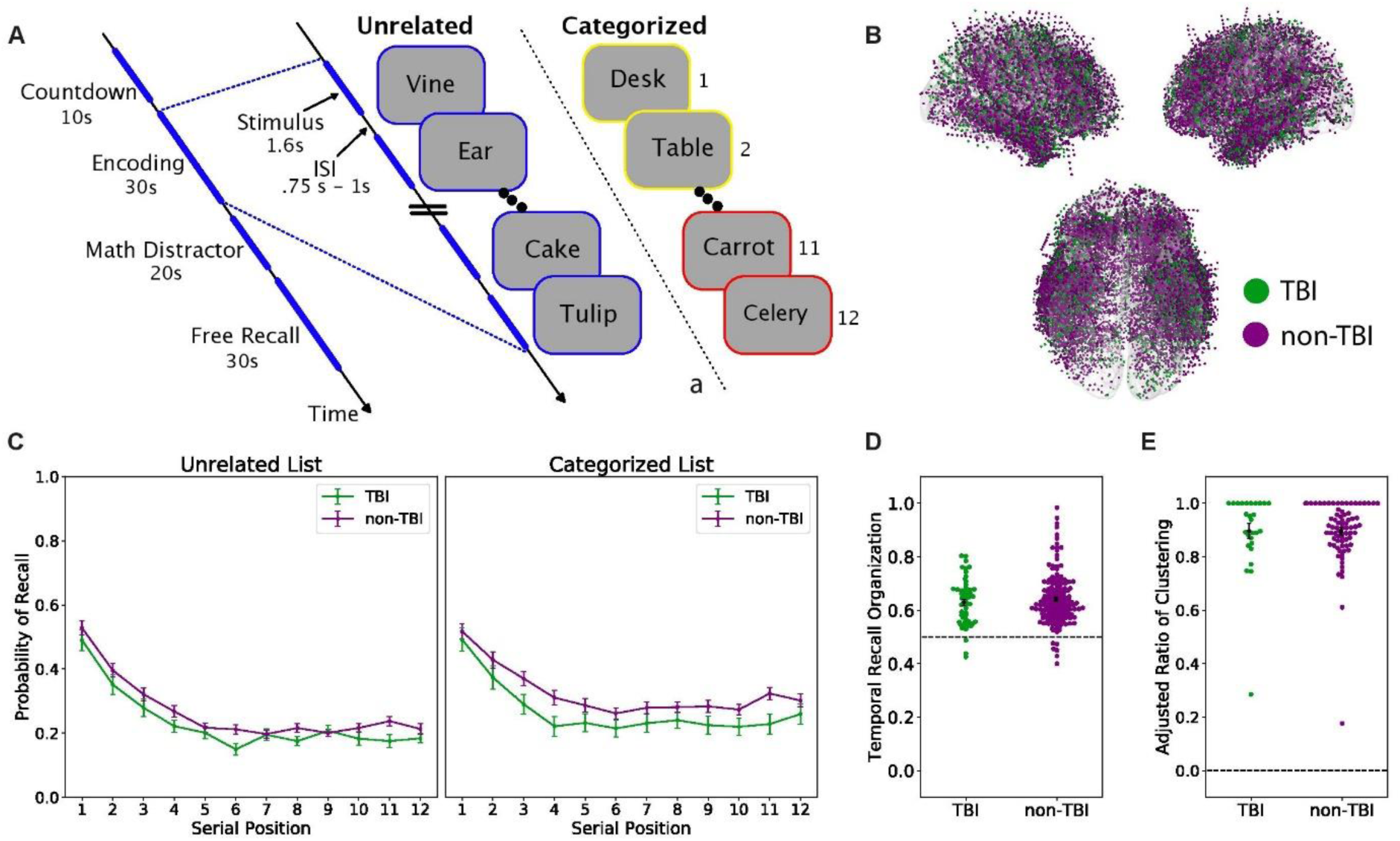
Memory task overview and performance. **(A)** Memory task schematic. In the delayed free recall task, subjects encoded a list of 12 items (words) presented sequentially, followed by an arithmetic distractor and verbal free recall, for up to 26 lists in a session. In a given session, all study lists contained either unrelated items, or items belonging to one of three semantic categories. **(B)** Localized intracranial electrodes overlaid on an average brain surface. Electrodes are colored by patient group (orange: TBI group, blue: non-TBI group) **(C)** Recall performance. Serial position curves plot the average probability of recall at each list position for the two groups. The TBI group trended towards lower recall performance compared to the non-TBI group (*p* = 0.074). The TBI group has significantly decreased recall performance in the categorized task (group *x* task interaction: *p* = 0.015) **(D)** Temporal clustering in recall. For each subject, we plot temporal clustering scores, with the group mean and standard error. The dotted line at 0.5 represents random chance. Both groups show temporal organization of their recall sequence, with no statistical difference between groups (*p* = 0.40). **(E)** Semantic clustering in recall. For each subject performing the categorized task, we plot the Adjusted Ratio of Clustering (ARC) score with group mean and standard error. Maximum clustering at 1 and no clustering at 0. Both groups exhibit semantic clustering of their recall sequences, with no significant difference between groups (*p* = 0.96).

Through careful review of medical records, a subset (*N* = 37) of patients (TBI group) were identified as having a history of moderate-to-severe TBI based on criteria developed by neurologists at the University of Pennsylvania (Dr. R. D-A.) and University of Texas Southwestern (Dr. K. D.), as follows: a reported history of significant head injury with either Loss of Consciousness (LOC) of at least 30 minutes or Post Traumatic Amnesia (PTA) of 24 hours or more, or a neuroradiology report describing lesions characteristic of TBI (Gupta *et al*., 2014). In this data-set, we identified 37 patients whose history indicated that they likely experienced a moderate-to-severe TBI. For 21 of these patients we obtained hospital records confirming all of the above criteria, for the remaining 17 patients, a careful review of hospital records conducted by two independent neurologists specializing in the treatment of TBI indicated a likely history of moderate-to-severe TBI although the hospital records lacked documentation confirming that the full set of criteria had been met. In **Table 1**, we provide demographics for the TBI group.

**Table 1:** TBI group demographics

|  | Age | Sex | Handedness | Age of Seizure Onset |
| --- | --- | --- | --- | --- |
| **TBI1** | 31 | F | R | 1 |
| **TBI2** | 39 | M | R | 16 |
| **TBI3** | 48 | M | R | 33 |
| **TBI4** | 34 | M | R | 1 |
| **TBI5** | 24 | M | R | 16 |
| **TBI6** | 35 | M | R | 15 |
| **TBI7** | 25 | M | R | 17 |
| **TBI8** | 57 | M | R | 6 |
| **TBI9** | 53 | M | R | 29 |
| **TBI10** | 28 | M | R | 12 |
| **TBI11** | 31 | F | R | 1 |
| **TBI12** | 38 | M | R | 33 |
| **TBI13** | 43 | F | R | 12 |
| **TBI14** | 44 | M | R | 17 |
| **TBI15** | 37 | M | R | 34 |
| **TBI16** | 47 | M | R | 44 |
| **TBI17** | 57 | M | R | 48 |
| **TBI18** | 33 | M | R | 13 |
| **TBI19** | 49 | M | L | 1 |
| **TBI20** | 36 | M | R | 2 |
| **TBI21** | 55 | M | R | 37 |
| **TBI22** | 29 | M | L | 4 |
| **TBI23** | 48 | M | A | 19 |
| **TBI24** | 60 | M | R | 50 |
| **TBI25** | 50 | M | R | 39 |
| **TBI26** | 55 | M | L | 20 |
| **TBI27** | 58 | F | A | 9 |
| **TBI28** | 45 | M | R | 18 |
| **TBI29** | 66 | F | L | 30 |
| **TBI30** | 51 | F | R | 11 |
| **TBI31** | 35 | M | R | 28 |
| **TBI32** | 54 | M | A | 39 |
| **TBI33** | 51 | M | R | 35 |
| **TBI34** | 47 | M | R | 22 |
| **TBI35** | 30 | M | R | 20 |
| **TBI36** | 32 | M | R | 27 |
| **TBI37** | 32 | F | R | 9 |

We identified the best matched control patients (non-TBI group) from the database (e.g. with no reported history of TBI) for each of the TBI subjects, by implementing a propensity score matching procedure (Rosenbaum and Rubin, 1983). First, we used a generalized linear model (GLM) to identify a set of relevant demographic and descriptive factors predictive of whether a patient was included in the TBI group: sex, handedness, prior resection, years of education, age at seizure onset, electrode coverage by brain region, and seizure onset zone by brain region. We constructed propensity score models (generalized linear models) of the above factors, with the regression estimating a similarity score for each non-TBI subject. We inputted the scores to the Matching package in R, “Multivariate and propensity score matching with balance optimization”(Sekhon, 2008), to determine the three best matches for each TBI subject, with exact matching required for handedness as an indicator of hemispheric dominance. This algorithm resulted in a 111 subject non-TBI group with characteristics that did not significantly differ from the TBI group, as highlighted in **Table 2**.

**Table 2:** Significant model covariates predictive of TBI

| | Age<br>(*Mean* ± *SD*) | Male<br>(%) | Right Frontal<br>Coverage (%) | Left Temporal<br>SOZ (%) | Right Temporal<br>SOZ (%) |
| --- | --- | --- | --- | --- | --- |
| **TBI (N=37)** | 42.9 ± 11.0 | 73.0 | 75.0 | 2.8 | 5.6 |
| **Match Group (N=111)** | 40.0 ± 11.5 | 70.3 | 73.3 | 2.9 | 5.7 |
| **All non-TBI (N=309)** | 36.4 ± 11.3 | 65.4 | 65.4 | 15.1 | 11.0 |

### Recall Tasks and Behavioral Analyses

Subjects completed between one and eight sessions (30-60 minutes duration) of a delayed free-recall task (Kahana, 2020). On each of up to 26 lists (per session), subjects studied 12 items, with each item appearing individually for 1.6 sec on a laptop display (a 0.75-1 sec inter-stimulus interval followed each item presentation). After the appearance of the last item, subjects performed a 20-second distractor task in which they solved a series of self-paced simple arithmetic problems. Next, a row of asterisks appeared, accompanied by a 1 second tone, to signal the start of a 30-second recall period, in which subjects freely recalled as many items as they could remember without regard to order of presentation.

Subjects participated in either, or both, of two variants of the free recall task described above, in which the studied items were either unrelated words or semantically-related (categorized) words. In the unrelated task, lists comprised of 12 common nouns drawn randomly and without replacement from a pool of 300 words chosen to have moderate-levels of memorability. For details on list construction, please see (Ezzyat *et al*., 2018). In categorized sessions, each 12-word list comprised four noun exemplars drawn from each of three taxonomic categories, with same category exemplars appearing in successively presented pairs such that the structure of a given list might be *A*_1_, *A*_2_, *B*_1_, *B*_2_, *C*_1_, *C*_2_, *B*_3_, *B*_4_, *A*_3_, *A*_4_, *C*_3_, *C*_4_. Pairs appeared in a random order constrained such that successive pairs always came from distinct categories. For details on category list composition see (Weidemann *et al*., 2019). As is customary in list memory experiments, we did not include data from the first (practice) list of each session in our subsequent analyses (Solway *et al*., 2012).

To assess the effect of prior injury history on recall performance, we used a linear mixed-effects model with group (TBI vs non-TBI), task, and serial position as fixed effects and random intercepts across subjects, and performed a likelihood ratio test, comparing the log-likelihood of the full model to a reduced model that did not include the main effect of group. To determine the significance of the interaction between group and task, we fit a second model that included interactions for all of the main effects, and compared it to a reduced model (without the group-task interaction) using the same likelihood ratio test described above. These analyses were programmed using the statsmodels library in python (Seabold and Perktold, 2010).

When freely recalling lists of items, the order of recalls reveals the dynamic, cue-dependent nature of memory retrieval. Sequentially recalled items exhibit several forms of “organization,” clustering according to their temporal and semantic relations (Kahana, 1996; Howard and Kahana, 2002). To quantify temporal organization, (Polyn *et al*., 2009) the temporal distance of each recall transition is compared to the distribution of temporal distances for all words that have not yet been recalled, in which 1.0 represents perfect temporal clustering and 0.5 represents chance. The average percentile score of all recalls for each subject is their temporal recall clustering score. For semantic clustering, we calculated an adjusted ratio of clustering (ARC) statistic (Roenker *et al*., 1971; Weidemann *et al*., 2019), in which a score of 1 represents maximum category clustering, and 0 represents clustering based on chance. For this analysis, we only included subjects from each group that completed the categorized variant of the task. For both metrics of recall, clustering scores for the two groups were statistically compared using a Welch’s independent t-test, which does not assume equal variance.

### Intracranial Recordings

We recorded EEG signals using varied clinical systems (Nihon-Kohden Neurofax EEG-1200, Grass Aura LTM64, XLTek EMU128FS, Natus Quantum) and research recording systems (Blackrock NeuroPort, Medtronic External Neural Stimulator) at sampling rates ranging from 250 to 2000 Hz. These data were either recorded at individual electrodes referenced to a common contact placed intracranially, on the scalp, or mastoid process and re-referenced to a bipolar referencing scheme or recorded directly using a bipolar referencing montage. In the bipolar scheme, neighboring contacts were referenced within each electrode. EEG data were aligned with behavioral data via TTL pulses sent from the behavioral computer to the EEG system, or through network packets passed between the behavioral computer and recording computer. The data were filtered using a fourth order Butterworth notch filter to remove 60 Hz line noise. The resulting signals were convolved with Morlet wavelets (wave number = 5) to obtain spectral power and phase measurements. As determined by a clinician, any contacts placed in epileptogenic tissue or exhibiting frequent inter-ictal spiking were excluded from all subsequent analyses.

To compare the activity in brain regions of interest, each electrode was localized to the patient’s specific anatomy by alignment and co-registration of pre-surgical T1 and T2 MRIs and post-implantation CT scans, including an automated segmentation of medial temporal lobe substructures (Yushkevich *et al*., 2014). Depth electrodes that were visible on CT scans were then localized within all the brain regions defined by the Desikan-Killiany Atlas (Desikan *et al*., 2006). Exposed recording contacts were approximately 1-2mm in diameter and 1-2.5mm in length; the smallest recording contacts used were 0.8mm in diameter and 1.4 mm in length.

### Analysis of spectral power and functional connectivity

To determine the relation between power and successful encoding in TBI and non-TBI subjects, we calculated the subsequent memory effect (SME) as the difference in spectral power (i.e. the power SME) for later recalled and not-recalled items. First, we calculated spectral power by convolving the raw EEG signal recorded during item presentation with 20 Morlet wavelets logarithmically spaced between 3 and 170 Hz. The resulting power values were log-transformed and z-scored within each session, and averaged over the 400-1100ms interval following item onset, resulting in a single power value for each item/frequency/electrode. We used a Welch’s t-test to quantify, for each electrode and frequency, the difference between later recalled and not recalled items. Here, a positive t-stat indicates an increase in power during successful memory encoding; a negative t-stat indicates a decrease. Next, we averaged these SME t-stats within four anatomical brain regions (Frontal, Medial Temporal, Lateral Temporal, Parietal) and two frequency bands: theta (4-8 Hz) and high frequency activity (HFA) (45-170 Hz). To assess differences in the average SME between the TBI and the non-TBI group, we used a Welch’s t-test with Benjamini-Hochberg FDR correction for multiple comparisons.

To assess differences in inter-regional connectivity in TBI and non-TBI subjects, we constructed a theta connectivity network for each subject by calculating the phase-locking-value, or PLV, (Lachaux *et al*., 1999) for every pair of electrodes. To compute the PLV, we extracted phase information during the 400-1100ms interval following item onset using 5 wavelets in the theta band (4, 5, 6, 7, and 8 Hz), calculated the average phase difference between each recording pair for every item, and measured the resultant vector (i.e. PLV) of phase differences for each item class (i.e. recalled vs not-recalled). We calculated the connectivity SME as the difference in PLV between these item classes, for each pair of electrodes and frequency in the theta band. A positive connectivity SME indicates greater synchronization at a given frequency between the pair of electrodes during successful memory encoding, and a negative SME represents greater synchronization during unsuccessful memory encoding. For each subject, we averaged the connectivity SME across all five frequencies to measure the overall theta-band connectivity SME, and pooled across all electrode pairs that spanned each unique region of interest (ROI) pair, yielding a theta connectivity SME for every ROI pair in the network.

To account for the imbalanced number of recalled and not-recalled items, we performed a non-parametric permutation test to compare the theta connectivity SME to a chance distribution. To generate a null (or chance) distribution of theta connectivity SME values for each electrode pair and subject, we randomly shuffled the item labels (i.e. recalled, not-recalled) 500 times and repeated the PLV computation for each shuffle. For each electrode pair, a z-score was derived by comparing the true theta connectivity SME value to its corresponding shuffled distribution. This z-score was taken to represent an edge weight of the connectivity network for each electrode pair in every subject, which were then averaged across electrode pairs that spanned ROI pairs. A positive z-score indicates an increase in synchronization between a pair of brain regions during successful encoding; a negative z-score indicates increased synchronization during unsuccessful encoding. To assess whether each group exhibited increased synchronization during successful memory encoding, we calculated an average edge weight across all ROI pairs for each subject, and compared the distribution of weights for each group to zero with a one-sample t-test. We used an independent t-test to compare the distributions of edge weights between the TBI and non-TBI groups.

### Multivariate Classification Methods

To discriminate patterns of spectral power predictive of successful memory encoding during the free recall task, we trained a logistic regression classifier using methods described in (Ezzyat *et al*., 2018). The input features were average spectral power for each item encoding epoch (0–1366 ms relative to item onset) calculated at each of eight frequencies log-spaced between 3 and 180 Hz and each electrode (frequency *x* electrode). The logistic regression model used L2-regularization with a penalty parameter (C) to 2.4*x*10^*−*4^ that was selected by a cross validation procedure on historical subjects via a grid search to correct for over-fitting. Given the possible class imbalance between later recalled and not recalled items depending upon subject performance, we weighted observations from the minority class in inverse proportion to the class frequency when training the classifier.

We quantified classifier performance by computing the average area under the curve (AUC), which measures the model’s ability to identify true positives while minimizing false positives, with chance *AUC* = 0.50. To mitigate the effects of overfitting on our evaluation of classifier performance, we used leave-one-out cross-validation when calculating AUC. For subjects performing multiple sessions of free recall, we used a leave-one-session-out (LOSO) method; for subjects with a single session, a leave-one-list-out (LOLO) method was used. To assess the significance of each subject’s individual classifier, we randomly permuted the recalled/not recalled labels in the training data and computed the AUC 1000 times to generate a null distribution. Classification analyses were programmed using the scikit-learn library in python (Pedregosa *et al*., 2011).

## Data Availability

De-identified data and analysis code is made available at: http://memory.psych.upenn.edu/Electrophysiological_Data

## Results

To support our analysis of the behavioral and physiological effects of TBI on memory biomarkers, we generated a matched group of controls over a set of relevant variables including age, sex, hemispheric dominance, seizure onset zone, and electrode coverage (see Methods). This yielded a non-TBI group of 111 subjects whose characteristics did not reliably differ from those of the 37 subject TBI group (see **Table 2**). **Figure 1A** illustrates our two memory paradigms: delayed free recall of (1) categorized and (2) unrelated lists of nouns. In both paradigms, subjects studied lists composed of 12 words for later recall, with each list followed by a brief arithmetic distractor task. Categorized lists comprised four exemplars drawn from each of three categories presented as successive pairs of categorized words. Unrelated lists were comprised of words drawn randomly and without replacement from a pool of common nouns (see Methods). As seen in **Figure 1B**, electrodes across both TBI and non-TBI subjects were distributed across similar regions, with the greatest density of electrodes sampling the temporal, medial temporal, and frontal regions.

We first asked whether the TBI and non-TBI groups differed in terms of their behavioral performance on our two memory tasks. Both groups exhibited typical serial position effects, recalling a higher proportion of early list items (the primacy effect) and a lower proportion of late-list items, which are attenuated in memory by the post-list arithmetic distractor task (see **Figure 1C** and Methods). Overall, TBI patients recalled fewer items than their non-TBI counterparts during both tasks, though this effect fell short of the significance threshold [*χ*^2^(1) = 3.2, *p* = 0.07]. Using a linear mixed-effects model, we found a significant interaction between task (categorized vs unrelated) and group (TBI vs non-TBI), in which the non-TBI group outperformed the TBI group to a greater extent during the categorized list paradigm [*χ*^2^(1) = 5.87, *p* = 0.015]. This finding is consistent with prior evidence of TBI-related impairment of semantic organization in verbal delayed free-recall (Vakil *et al*., 2019). Further, analysis of both groups’ recall dynamics revealed significant temporal and semantic clustering, recapitulating well-characterized episodic memory dynamics. List items were more likely to be successively recalled if they were studied in nearby list positions [Temporal: TBI *t*(51) = 10.6, *p <* .01; non-TBI *t*(166) = 18.2, *p <* .01] (**Figure 1D**) and came from the same category [Semantic: TBI *t*(28) = 31.6, *p <* .01; non-TBI *t*(75) = 66.2, *p <* .01]. There was no significant difference between the two groups for these measures of recall organization [Temporal: *t*(217) = −0.84, *p* = 0.40; Semantic: *t*(103) = 0.04, *p* = 0.96] (**Figure 1E**).

To determine whether the neural activity underlying successful memory encoding differed between TBI and non-TBI cohorts, we employed the subsequent memory effect (SME) paradigm. The SME compares neural measures during encoding of items that were subsequently recalled with items that were subsequently not recalled, reflecting activity that correlates with memory success. As prior work has established that increased high-frequency activity (HFA, 45-170 Hz) and increased inter-regional theta synchrony mark periods of successful memory encoding (Burke *et al*., 2014; Solomon *et al*., 2017), we conducted this subsequent memory contrast for two key biomarkers of successful memory formation: spectral power and theta (4-8 Hz) phase connectivity, asking whether these particular biomarkers serve as indices of successful memory encoding in subjects with a history of moderate-to-severe TBI.

We first compared patterns of whole-brain spectral power SMEs between the TBI and non-TBI groups. To do this, we computed spectral power at a broad range of frequencies (3-170 Hz) during the item encoding period, for each electrode. We then statistically compared the distribution of powers for items that were later recalled to powers for items that were not recalled, generating a spectral power SME t-stat (see Methods). **Figure 2A** illustrates the average power SME t-stat at 20 different frequencies and 4 brain regions associated with episodic memory (frontal lobe, medial temporal lobe (MTL), lateral temporal cortex (LTC), parietal lobe) in the two patient subgroups. As in prior studies, we found a significant increase in high frequency activity [*p<*.05]; Benjamini-Hochberg corrected] in both the TBI and non-TBI groups in all regions except the LTC. We further noted a significant decrease in low frequency activity [*p <* .05; Benjamini-Hochberg corrected] in all four brain regions in both groups. There were no significant differences in SMEs between the two groups at any brain region in the theta and HFA frequency bands [*p >* 0.1] as seen in **Figure 2B**.

**Figure 2.**
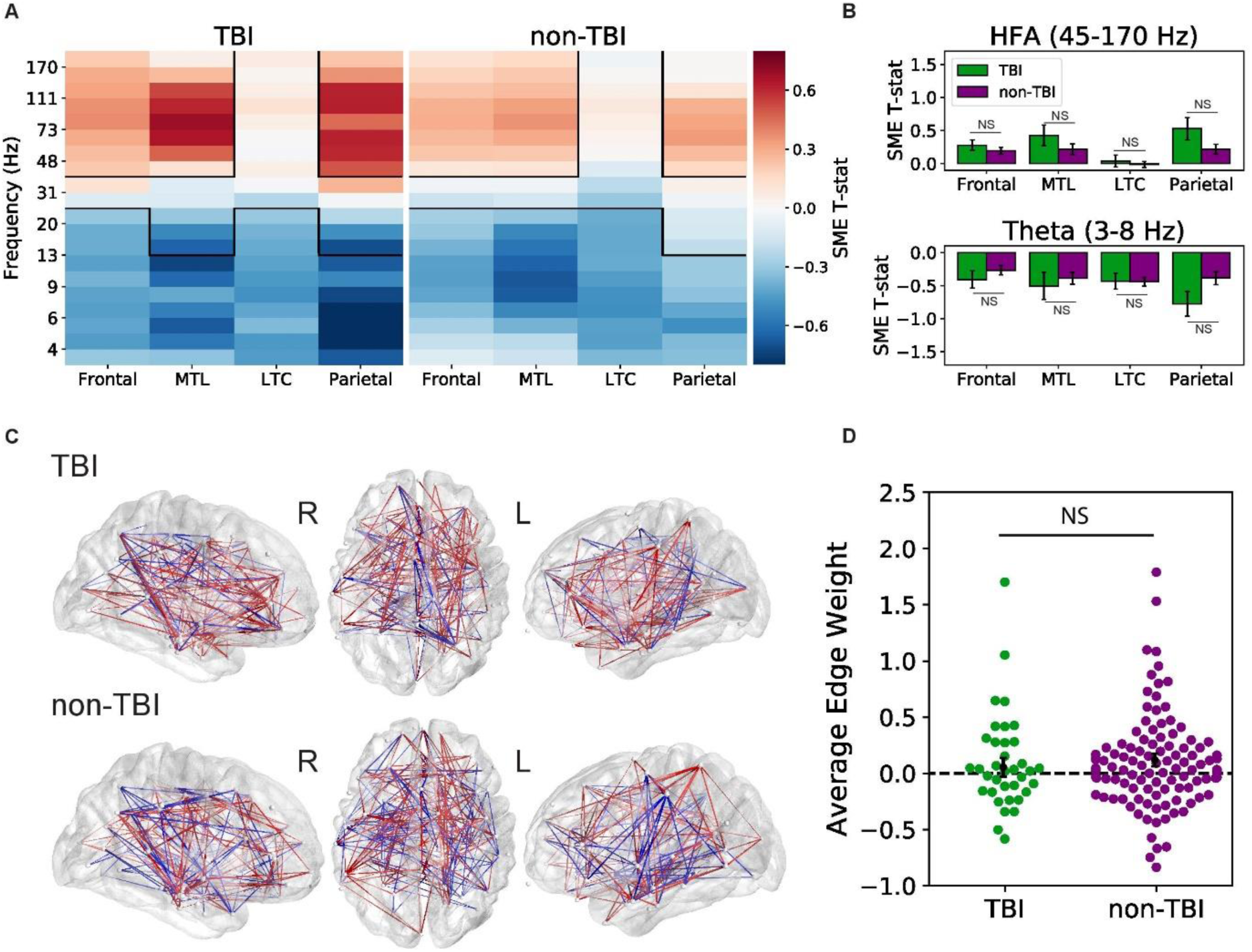
Biomarkers of successful memory encoding. **(A)** Average spectral power SME t-stats (recalled vs. non-recalled encoding periods) at 20 frequencies spanning 3 to 170 Hz and 4 distinct brain regions (Frontal lobe, Medial temporal lobe (MTL), Lateral Temporal Cortex (LTC), and Parietal lobe). Both groups exhibit a significant increase in high frequency activity (HFA), a significant decrease in low frequency activity during successful encoding [*p <* 0.05], **(B)** and statistically similar power SME in every ROI [*p >* 0.1] **(C)** Group average theta encoding networks overlaid on an average brain surface. Each node represents a distinct ROI. Connections (edges) between nodes are colored red when theta synchronization increases during successful memory encoding and blue where synchronization decreases during encoding. **(D)** Average synchronization z-score (‘edge weight”) distributions, with positive values reflecting overall memory-related synchronization. Across subjects, we found a significant increase in theta synchronization with successful memory encoding for the non-TBI group [*p* = 0.03] but not for the TBI group [*p* = 0.51]; there was no evidence of a difference between the two groups [*p* = 0.52].

To assess for differences in functional networks underlying successful memory encoding between TBI and non-TBI patients, we analyzed patterns of inter-regional phase connectivity in the theta band. To construct networks of connectivity between electrodes in each subject’s brain, we calculated the phase-locking value (or PLV), which quantifies the phase synchronization in the theta band (4 to 8 Hz) for every ROI pair. We calculated a connectivity SME z-score, reflecting the difference in PLV between successful and unsuccessful item encoding events; a positive value indicates increased phase synchrony between ROIs during successful encoding (see Methods). Connections with Z-scores greater than 1.96 are displayed in **Figure 2C**, where red lines represent memory-related increases in connectivity, while blue lines represent memory-related decreases in connectivity. Qualitatively, these maps show strong fronto-temporal synchrony during successful memory encoding in both TBI and non-TBI groups.

To determine if these theta connectivity networks reliably differed between TBI and non-TBI groups, we took the average PLV z-score across all connections (or “edge weights”) in each subject as a measure of overall synchronization during good memory states. Congruent with previous findings, both the TBI and non-TBI subject groups showed a general increase in theta synchronization during successful encoding; average z-scores were above 0 in both groups, indicating an overall propensity for memory-related synchronization. As seen in **Figure 2D** the non-TBI group had z-score averages that were significantly higher than zero [*t*(110) = 2.20, *p* = 0.03], though the TBI group did not meet significance [*t*(36) = 0.66, *p* = 0.51]. However, we did not find a significant difference in the average z-score between the two groups [*t*(146) = *−*0.65, *p* = 0.52].

The ability to reliably predict subsequent memory in real-time is potentially a critical tool for designing new therapies and interventions. Here, we trained an individual multivariate logistic regression classifier for each subject to discriminate patterns of spectral power during memory encoding, effectively forecasting whether each item would be later recalled or forgotten. In **Figure 3**, we plot the average true positive and false positive rate for probability thresholds ranging from 0 to 1 as receiver operating characteristic (ROC) curves. To quantify classifier performance, we computed the average area under the curve (AUC) for each subject’s classifier and averaged within each group. Overall, the TBI group had an average AUC of 0.624 [*SEM* = 0.011], and the non-TBI group had an average AUC of 0.620 [*SEM* = 0.008] with statistically equivalent distributions [*t*(142) = 0.29, *p* = 0.77]. This result suggests that spectral power features of recorded intracranial EEG can effectively predict memory performance in both TBI and non-TBI patients, and could be used to responsively trigger closed-loop brain stimulation.

**Figure 3.**
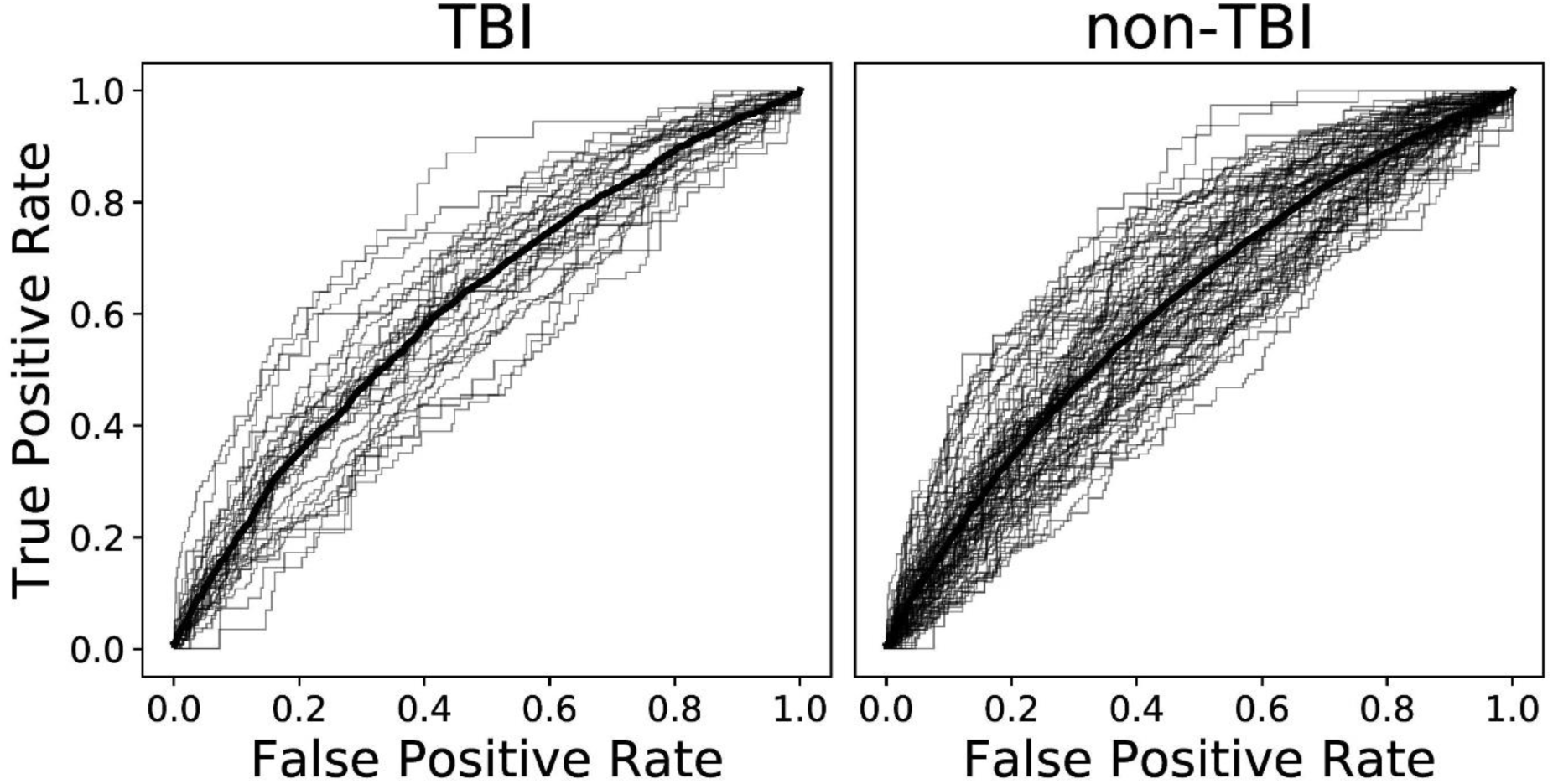
Classifier performance. For each unique subject, receiver operating characteristic (ROC) curves show the performance of a logistic regression classifier tested on held-out sessions of the delayed free recall task (grey lines), with the group average performance shown in bold (TBI group, left: mean *AUC* = 0.624, *N* = 37; non-TBI group, right: mean *AUC* = 0.620, *N* = 111).

## Discussion

Our mnemonic ability exhibits marked variability from moment to moment and from day to day, independent of item characteristics and task variables (Kahana *et al*., 2018).

Measures of brain activity similarly reflect this behavioral variation, constituting neural biomarkers of mnemonic success. Direct brain recordings from neurosurgical patients have revealed two striking biomarkers of effective memory encoding: First, the power spectrum of intracranial EEG activity tilts upward during periods of successful encoding. Second, theta frequency neural activity (3-10 Hz) becomes coherent and synchronous during periods of successful encoding (Burke *et al*., 2013; Solomon *et al*., 2017). These two patterns emerge across a broad network of brain regions, described as the” corememory network” (Wagner, 1998; Paller and Wagner, 2002; Kim, 2011; Burke *et al*., 2014; Long and Kahana, 2015). Here we examined these two biomarkers of successful memory function in matched groups of neurosurgical patients with and without a significant prior-history of traumatic brain injury.

Traumatic brain injury produces diverse and heterogeneous damage to neural networks including diffuse axonal injury, diffuse microvascular injury, and neuroinflammation, and ultimately, impaired memory ability in many patients. As such, we wondered whether patients who experienced moderate-to-severe TBIs would exhibit differences in their biomarkers of successful memory function.

To answer this question, we recorded from indwelling electrodes in cohorts of TBI-affected and non-TBI neurosurgical patients as they performed verbal free-recall tasks. Although our TBI group exhibited marginally worse recall performance, both groups showed nearly identical serial position effects, temporal and semantic clustering (see **Figure 1C**,**D**,**E**). Both groups also exhibited highly similar electrophysiological correlates of successful memory (**Figure 2**), suggesting that biomarkers of local memory processing were unaffected by TBI neuropathology. Finally, we examined the utility of spectral power biomarkers of successful memory in reliably predicting recall performance. We found that this approach yielded statistically equivalent classification ability for both the TBI and non-TBI groups, with both groups showing statistically reliable classification (AUCs of 0.0624 and 0.0620 for TBI and non-TBI groups, respectively).

The surgical treatment of refractory epilepsy occasionally requires long-term implantation of electrodes for seizure localization. By generously volunteering to take part in cognitive studies, patients undergoing such treatment have afforded scientists an unprecedented view of the neural basis of human cognition. Whereas only a handful of epilepsy centers conducted this type of cognitive electrophysiological research at the dawn of the 21st century, we are now seeing scientific reports from most of the major epilepsy centers worldwide. With this advancing research program characterizing the neural correlates of diverse cognitive processes, the neuroscientific community must consider whether these findings generalize beyond the population of epilepsy patients. This question of generalization has obvious scientific and therapeutic implications.

Here we sought to determine whether spectral correlates of successful memory, studied and extensively documented in epilepsy patients (Ezzyat *et al*., 2017; Herweg *et al*., 2020), generalize to those patients with an additional history of TBI, or if TBI-related pathology produces distinct biomarkers of variability in memory function. In conducting this study, we did not know what to expect. Given that a history of TBI suggests an underlying pathology which differs from that seen in non-TBI epilepsy patients (Bigler and Maxwell, 2011), we did not expect to find such striking similarities among the two groups.

The striking similarity between biomarkers of successful memory in TBI and non-TBI groups helps to address a fundamental question in human neuroscience; namely, do findings in patients with epilepsy-related pathology generalize to other clinical subgroups?

The invariance of biomarkers across clinical subgroups bolsters arguments that task-manipulation contrasts identify neural processes that more likely reflect normal brain function (Kahana *et al*., 1999; Kahana *et al*., 2001). That is, although TBI and epilepsy-related pathology both disrupt neural mechanisms that support memory, these mechanisms still function to some extent, and comparisons of successful and unsuccessful items help to identify these residual functions. Of course, this hypothesis need not be true, and future work across more diverse patient populations may well temper the claims laid out in this paper.

Although our study can only speak to the similarity of a particular set of biomarkers of memory across two distinct epilepsy subgroups, this particular inter-group comparison has important clinical implications. Recent studies have shown that closed-loop neuromodulation in refractory epilepsy patients can reliably boost memory, albeit to a modest degree (Ezzyat *et al*., 2018), and closed-loop DBS promises to improve existing therapies in other patient populations, such as Parkinson’s (Swann *et al*., 2018). Such findings raise hope for the use of closed-loop electrical stimulation as a therapy for memory loss in patients with brain injury or other types of neurological or neurogenerative diseases. Our findings suggest that biomarkers used to guide such therapy appear conserved in at least some epilepsy patients with an additional history of traumatic brain injury, paving the way for future tests of closed-loop neuromodulation in TBI patients.

## Acknowledgements

We thank Blackrock Microsystems and Medtronic for providing neural recording equipment. We owe a special thanks to the patients and their families for their selfless participation and support of the study. We thank Nora Herweg for her valuable feedback on this work.

